# Parkinsonian gait effects with DBS are associated with pallido-peduncular axis activation

**DOI:** 10.1101/2021.10.13.464270

**Authors:** Mojgan Goftari, Chiahao Lu, Megan Schmidt, Remi Patriat, Tara Palnitkar, Jiwon Kim, Noam Harel, Matthew D. Johnson, Scott E. Cooper

## Abstract

**Background:** Deep brain stimulation (DBS) targeting the subthalamic nucleus (STN) often shows variable outcomes on treating gait dysfunction in Parkinson’s disease (PD). Such variability may stem from which specific neuronal pathways are modulated by DBS and the extent to which those pathways are modulated relative to one another.

**Objective:** Leveraging ultra-high-field (7T) imaging data and subject-specific computational models, this study investigated how activation of seven distinct pathways in and around STN, including the pallidopeduncular and pedunculopallidal pathways, affect step length at clinically-optimized STN-DBS settings.

**Methods:** Personalized computational models were developed for 10 subjects with a clinical diagnosis of PD and with bilateral STN-DBS implants.

**Results:** The subject-specific pathway activation models showed a significant positive association between activation of the pedunculopallidal pathway and increased step length, and negative association on step length with pallidopeduncular pathway and hyperdirect pathway activation.

**Conclusions:** The STN region includes multiple pathways, including fibers of passage to and from the mesencephalic locomotor area. Future clinical optimization of STN-DBS should consider these fibers of passage in the context of treating parkinsonian gait.

## Introduction

Parkinsonian gait dysfunction, which is characterized in part by shortened step length, is difficult to treat consistently with STN-DBS therapy^1–4^. This variability likely stems from multiple factors. First, it is difficult to optimize stimulation settings for gait within a clinical visit because DBS has relatively long wash-in and wash-out time constants for those symptoms^5^. In some cases, DBS can also worsen gait, especially when stimulating through electrodes in regions dorsomedial to the STN^6,7^. Finally, robust therapy for parkinsonian gait may require direct modulation of motor control networks beyond the classic basal ganglia circuit^8,9^.

The STN region is a nexus of motor control circuits that project between the brainstem, basal ganglia, thalamus, and cortex. While pathway activation models of STN-DBS have focused on axonal projections between the external globus pallidus (GPe) and STN^10–14^, from the motor cortices to STN (hyperdirect pathway)^11,15–17^, and from the internal globus pallidus (GPi) coursing around STN and to the motor thalamus^10,13,16^, many other pathways are likely modulated by STN-DBS. For example, primate neuroanatomical tracing studies have shown reciprocal axonal connections between the pedunculopontine nucleus (PPN) and the basal ganglia, with axons coursing through the STN (pedunculopallidal tract^18^) or around the STN (pallidopeduncular tract^19^). These tracts when stimulated may engage the mesencephalic locomotor region^20^ differentially through antidromic and orthodromic activation, respectively^21^. Indeed, clinical studies in PD patients have suggested that direct stimulation of pallidofugal efferents through GPi-DBS can exacerbate axial symptoms^22^ and in non-PD patients can induce hypokinetic gait, decreased step length, cadence increase, and freezing of gait^23,24^.

Computational pathway activation models of DBS are useful in assessing the degree to which brain circuits are activated by stimulation settings that do or do not improve clinical symptoms. Knowledge gained in this way can be leveraged by optimization algorithms to reduce the number of possible stimulation settings to be evaluated clinically^12,25^. To this end, in ten individuals with bilaterally implanted STN leads, we developed subject-specific models of seven distinct pathways in and around the STN incorporating two new pathways, pallidopeduncular and pedunculopallidal, in order to test our prospective hypothesis that targeting axons within the pallidopeduncular pathway would decrease step length while activation of the pedunculopallidal pathway would increase step length.

## Methods

### Human subjects

Ten individuals with a diagnosis of Parkinson’s disease and bilateral STN-DBS were recruited (**Table 1**). Prior to STN-DBS lead implantation, subjects received a ultra-high-field (7T) brain MRI (7T MAGNETOM 90cm bore scanner) at the Center for Magnetic Resonance Research (CMRR) at the University of Minnesota^26–28^. Approximately one month post-implant, subjects underwent head CT (0.6 mm slices). DBS leads included (1) a 4-channel lead with ring electrodes (Medtronic, 3389 lead) in 4 subjects, (2) an 8-channel lead with ring electrodes (Boston Scientific, 2201 lead) in 3 subjects, and (3) an 8-channel directional lead with two split-band rows between ring electrodes (Abbott, 6172/6173 leads) or between a ring electrode above, and cap electrode below (Boston Scientific, 2202 lead) for 3 subjects. Following implantation, stimulator “programming” adjustments were made over several months by a subspecialized clinician independent of this study. All study participants had reached stable stimulator settings by the time gait was tested for this study. All procedures were approved by the Institutional Review Board of the University of Minnesota, and subjects gave their written consent to participate in the study.

**TABLE 1.** Clinical features and DBS settings of subjects in this study

| Subject | Sex | Age | UPDRS III<br>(Off-Med) | UPDRS III<br>(On-Med) | PD<br>duration<br>(years) | Implant<br>duration<br>(years) | DBS<br>Lead<br>Model | Clinical DBS<br>Setting (R,L) |
| --- | --- | --- | --- | --- | --- | --- | --- | --- |
| 1 | M | 66 | 41 | 28 | 11 | R:2, L:3 | BSC 2201 | C(+)E3(-)E6(-), C(+)E3(-) |
| 2 | M | 84 | 18 | 11 | 8 | R:6, L:2 | MDT 3389 | C(+)E2(-), C(+)E3(-) |
| 3 | M | 73 | 42 | 24 | 15 | R:6, L:7 | MDT 3389 | C(+)E2(-), C(+)E2(-) |
| 4 | M | 74 | 31 | 17 | 14 | R:5, L:5 | MDT 3389 | C(+)E2(-), C(+)E2(-) |
| 5 | M | 61 | 39 | 7 | 17 | R:4, L:4 | BSC 2201 | C(+)E2(-), C(+)E3(-) |
| 6 | F | 68 | 36 | 17 | 7 | R:1, L:1 | BSC 2202 | C(+)E2b(-), C(+)E3c(-) |
| 7 | M | 69 | 40 | 8 | 14 | R:3, L:3 | ABT 6172 | C(+)E3(-), C(+)E2(-) |
| 8 | F | 68 | 20 | 7 | 11 | R:2, L:1 | ABT<br>6172/6173 | ILS C(+)E3c(-) & E2(+)E4(-),<br>C(+)E3b(-) |
| 9 | M | 62 | 42 | 15 | 16 | R:6, L:6 | MDT 3389 | C(+)E1(-), C(+)E2(-) |
| 10 | M | 64 | 27 | 11 | 10 | R:4, L:4 | BSC 2201 | E3(+)E2(-), E1(+)E2(-) |
| Sum | 2F/8M | 68.9±6.4 |  |  |  |  |  |  |

MDT: Medtronic, BSC: Boston Scientific, ABT: Abbott
\*E1 is the bottom electrode for the Boston Scientific and Abbott lead
\*E0 is the bottom electrode for Medtronic lead

### Clinical assessment of parkinsonian gait

Subjects started a gait testing session in the practically-defined off medication state (Off-Med), having taken no regular PD medications for 12 hours and no daily PD medications (rasagiline, Mirapex ER, Requip XL) for 24 hours. They arrived with their neurostimulators turned on, at their usual, clinical DBS settings. In this state, they first walked, at an instructed “ordinary, comfortable speed” for 20 feet in a hallway, timed with a stopwatch, in order to determine natural overground speed (average of 6 trials).

Subjects next began walking on an instrumented treadmill (CMill, Motek-Hocoma). We chose treadmill walking for our measurements, because it allowed for a large number of gait cycles in a compact space. Three treadmill speeds were used: 100%, 85%, and 75% of each subject’s natural overground speed. While walking, patients wore a safety harness attached to an overhead sliding track that allowed unrestricted movement, while protecting against the possibility of a fall. The treadmill’s tachometer measured actual belt speed, and its integral forceplate measured the center of pressure (COP) from the subject’s feet. From the cyclical COP trajectory, the manufacturer’s software computed the time and location of the left and right heel-strike and toe-off. These, and the belt speed, were exported and processed to compute left and right step lengths (left heel-strike to right heel-strike, and vice versa). These measurements have been validated against overground gait measurements obtained with a pressure-sensitive mat (Gait Rite, CIR Systems)^29,30^.

Treadmill walking consisted of 1 minute trials at steady-state (plus a few seconds at the beginning and end while the treadmill ramped up to speed, and back down to stationary to avoid unbalancing the subject). One minute of seated rest preceded each trial. Treadmill testing began with a practice trial at 100% of overground speed followed by trials at 75%, 85%, and 100% speed, which formed the baseline data for the “wash-out” phase of the experiment. DBS was then turned off, and the subject immediately did three more trials at 75%, 85%, and 100%. Sets of trials at these three speeds were then repeated every 15 minutes for the next hour, for a total of 5 sets, the last occurring one hour after DBS was turned off. Following a rest break, another three trials at 75%, 85% and 100% were collected as the baseline data for the “wash-in” phase of the experiments. DBS was then turned on, and the subject immediately did a set of three more trials at those speeds. Sets of trials at these three speeds were then repeated every 15 minutes for the next hour, for a total of 5 sets, the last occurring one hour after DBS was turned on. One subject completed only 4 sets of wash-in trials due to urinary urgency.

### Subject-specific anatomical reconstructions and pathway models

Pre-operative anatomical T1-weighted (7T, 0.6 mm isotropic), T2-weighted (0.4×0.4×1 mm), susceptibility-weighted imaging (SWI; 0.4×0.4×0.8 mm) and diffusion-weighted MR imaging data (1.5 mm isotropic) were used to construct subject-specific models. Diffusion-weighted data were processed for motion, susceptibility and eddy currents distortions correction using the FDT diffusion and topup tools in FSL (FMRIB Software Library v6.0). Subject-specific neural pathway models were generated from a combination of probabilistic fiber tractography and visualization and segmentation of the pathways using the subject’s own ultra-high-field 7T imaging data. Modeled pathways included the: (1) corticospinal tract of internal capsule (cstIC), (2) cortico-subthalamic tract (hyperdirect pathway, HDP), (3-5) full morphological reconstructions of STN neurons parcellated into overlapping functional territories, (6) pedunculopallidal tract, and (7) pallidopeduncular tract. To model the cstIC, probabilistic fiber tractography was performed on the subject-specific DTI data using FSL. As previously published^15,12^, seed points were placed in the primary motor cortex based on segmentation using FreeSurfer’s cortical parcellation of precentral gyrus (Destrieux atlas) and ipsilateral crus cerebri (T1-weighted data). Additionally, the HDP was modeled as a set of collaterals emerging from cstIC and terminating at random points in the segmented volume of the STN^12,15,17^ (**Fig. 1A**).

**Figure 1.**
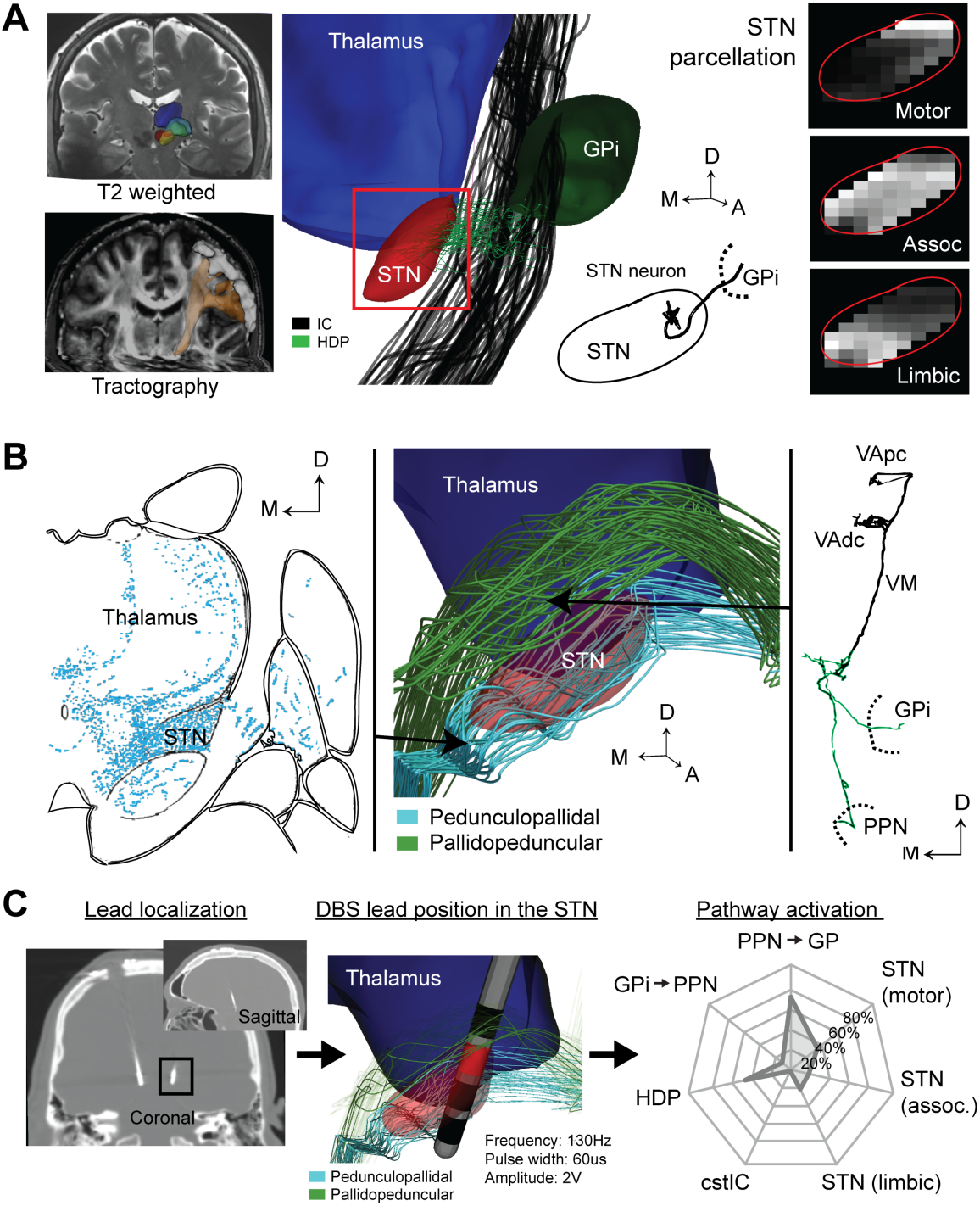
Personalized, subject-specific pathway activation models. **(A, B)** Seven anatomical pathways were digitally reconstructed from each subject’s ultra-high-field 7T MRI data. These pathways included subthalamopallidal neurons from motor, associative, and limbic probabilistic parcellations of STN; hyperdirect pathway; cortico-spinal tract of internal capsule; pedunculopallidal pathway; and pallidopeduncular pathway. **(C)** Localization of each subject’s DBS lead placement enabled calculating biophysical activation profiles for each neural pathway based on the DBS settings tested within the clinic. M: medial; D: dorsal; A: anterior. Other acronyms are provided in the text.

The STN was manually segmented using a combination of T1-weighted, T2-weighted and SWI imaging data. STN was then parcellated into motor, associative, and limbic territories using a tractography-based method^27,28^ (**Fig. 1A**). Briefly, probabilistic tractography was generated with STN as a seed and the ipsilateral cortical regions of the brain (motor, associative, limbic) as target masks and contralateral cortical regions as exclusion masks. Each STN voxel was considered to be connected to a cortical region if at least 25% of the probabilistic tracts connected the two regions. This allowed for one voxel to be connected to more than one cortical area. STN neurons were modeled such that they had five to eight long, sparsely spiny dendrites that were arborized mostly along the principal axis of the nucleus. These neurons had one axonal branch coursing rostrally and laterally toward the globus pallidus^31,32,10^.

The pedunculopallidal and pallidopeduncular pathways were modeled using segmentations of the 7T T1 imaging data and axonal trajectories based on previous primate histological tracing studies^18,19^. Ascending efferents from the pedunculopontine nucleus (PPN) were modeled as a broad axonal tract coursing through the STN en route to the globus pallidus via the ansa lenticularis^18^. Globus pallidus internus (GPi) efferents were modeled as coursing immediately dorsal to and around the anterior two-thirds of the STN and then projecting through the lenticular fasciculus and caudal to the PPN (H2, pallidopeduncular tract) (**Fig. 1B**).

### Modeling effects of electrical stimulation

The trajectory of each DBS lead and orientation of individual electrode contacts were determined from CT scans using the unique artifact characteristics of the segments and fiducial marker^33^. Post-operative CT scans were co-registered to the pre-operative MRI using Elastix^34,35^. The extracellular voltage distribution resulting from the clinically applied DBS configuration was calculated using an inhomogeneous and anisotropic finite element model specific to the anatomy of each subject. In the NEURON programming environment^36^, the subject-specific stimulus waveform was applied to each compartment of the biophysical pathway models^12–14^ to predict transmembrane currents through the modeled neurons or axons. An axon was considered ‘activated’ if an electrical pulse train generated phase-locked action potentials that propagated to the distal end of the modeled axon (**Fig. 1C**). The percentage of axons within a pathway activated by each DBS configuration was calculated, which defined a pathway activation measure. This approach differs from previous studies using the volume of tissue activated (VTA), which is known to be less accurate and less informative than the pathway-specific approach that we used in this study^11^.

### Statistical analysis

We investigated the effect of the neural pathway activations computed by the computational models on changes in step length between on-DBS and off-DBS conditions. We used MATLAB 2018a and the *fitlme* function to construct a nested repeated measures random intercept model with *side* in *subject* as random effect and percent pathway activations as fixed factors. The methods used for approximation of the coefficients and confidence intervals were Maximum Likelihood and Residual method, respectively. These statistical methods were justified because the distribution skewness of the percent change in step length was normal and balanced^37^ (Pearson 2 skewness coefficient=0.2973, n=40). The model resulted in a coefficient estimate and a confidence interval for each fixed term. A large positive value of the coefficient estimate signified a strong positive correlation between independent variable (e.g., pedunculopallidal pathway activation) and dependent variable (percent change in step length). Data were analyzed to test the hypothesis that PPN→GP activation will increase step length whereas GPi→PPN activation will decrease step length; additionally, exploratory analysis was performed on how other pathway activations related to changes in step length with DBS. Statistical significance was assessed using an F-test at *p*<0.05.

## Results

### Changes in step length across subjects

Contralateral step length (averaged over each one-minute trial) was expressed as a percent change relative to baseline at the corresponding speed (75%, 85% and 100% of overground speed) and tabulated for all 10 subjects (20 brain hemispheres) at each time point, during wash-in and wash-out portion of each experiment (filled circles in **Fig. 2A**). The change in contralateral step length generally stabilized within 30 minutes for both wash-in and wash-out. To account for these dynamics, we fit an exponential curve with two time constants [i.e. exp(-a_1_t) +exp(-a_2_t)] to the repeated measures (dashed lines in **Fig. 2A**) and calculated the step length changes at steady-state (unfilled circles in **Fig. 2A**). Steady-state step length changes ranged from -30% to 20% (**Fig. 2A**, with a positive value indicating an increase in step length) amongst the 20 individual DBS leads. Comparing the steady-state step length changes during wash-in (on-DBS) conditions showed step length increased for 60% of the DBS leads evaluated (**Fig. 2B**). As expected, there was also an inverse relationship between the changes in step length during wash-in and the changes during wash-out (off-DBS).

**Figure 2.**
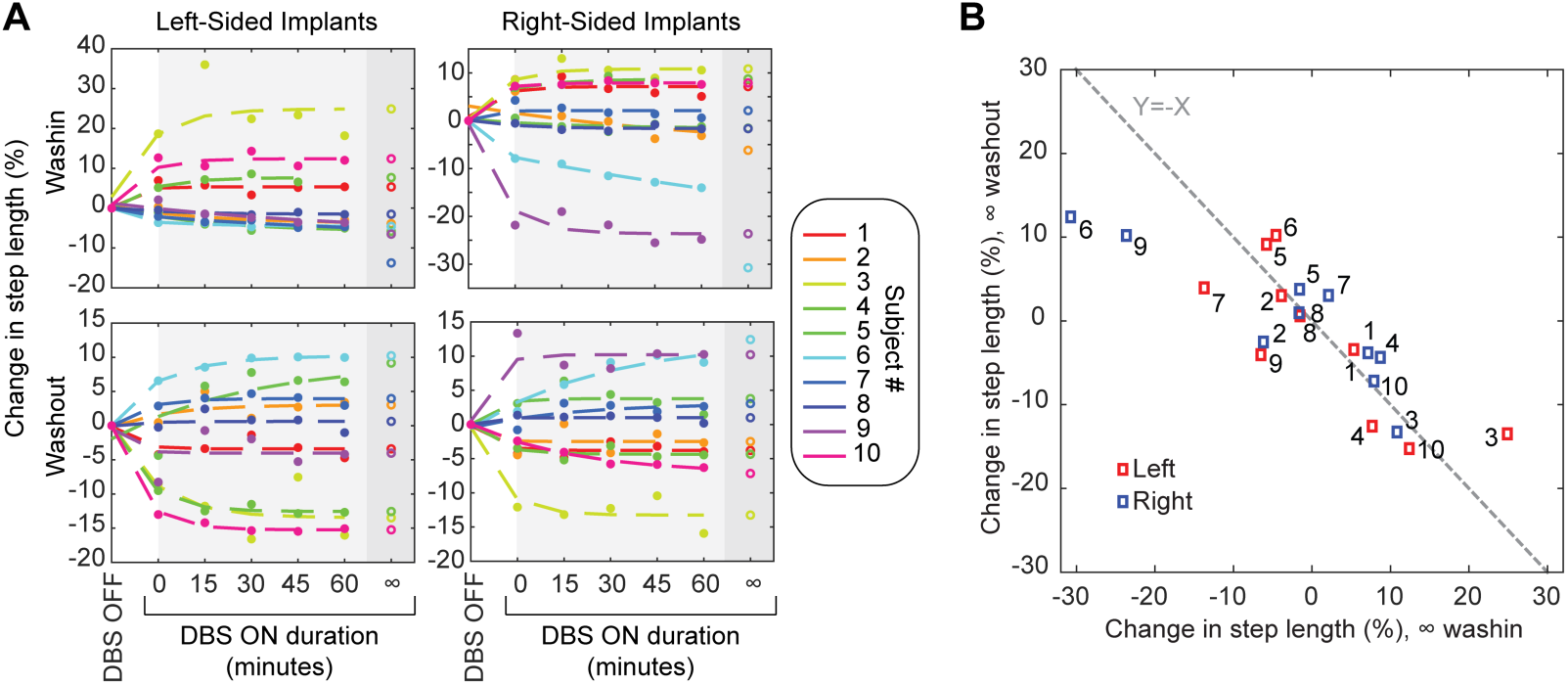
DBS-induced change in step length during wash-in and wash-out conditions. **(A)** Clinical step length changes (filled circles) followed an exponential model with two time constants. Unfilled circles show the asymptotes of the fit. **(B)** The modeled step length changes were similar in magnitude and opposite in polarity between wash-in and wash-out.

### Degree of pathway activation in and around the STN at clinically optimized settings

Across the twenty DBS leads studied, the pathway activation models showed that STN-DBS activated a distributed set of pathways within and adjacent to the STN that were specific to each subject (**Fig. 3**). Amongst subjects, the pedunculopallidal (PPN→GP) pathway had the strongest mean activation (92.0%), followed by the GPi→PPN pathway (73.9%), the hyperdirect pathway (63.2%), and the subthalamic nucleus efferent pathways (motor: 28.2%, associative 27.2%, and limbic 22.8%). The mean percent activation for the STN motor pathway was higher than that for STN associative and limbic pathways, which was expected since the STN-DBS lead implants most often traversed the posterolateral portion of the STN in this cohort of subjects (**Fig. S1**). In the context of interpreting associations between pathway activations and symptoms, it is important to consider the correlation amongst pathway activations (**Fig. S2A)**. There were strong correlations amongst activations for the STN motor, STN associative, and STN limbic pathways. Other pathways showed weaker correlations in pathway activation amongst the leads (**Fig. S2B**).

**Figure 3.**
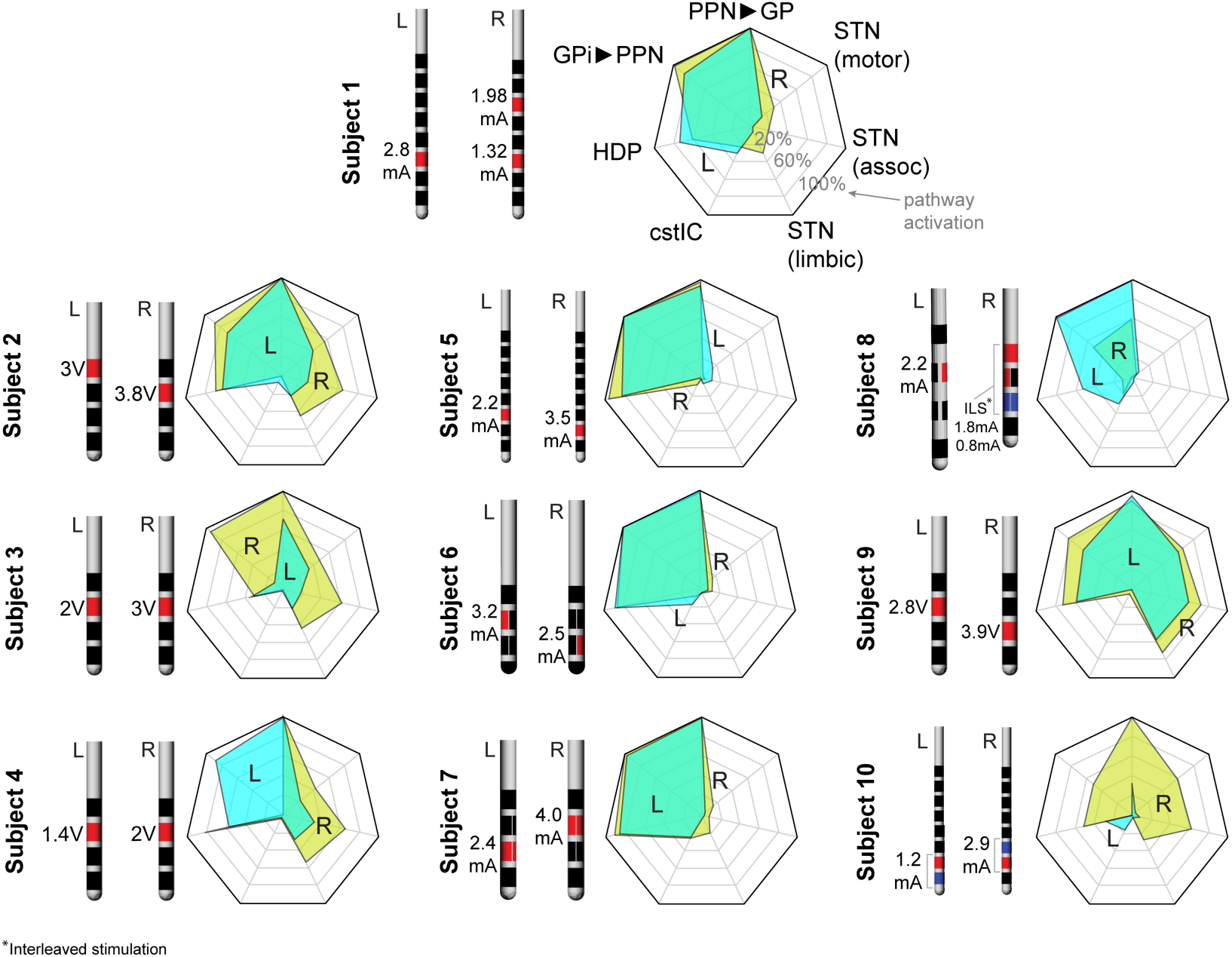
Pathway activation maps for each DBS lead tested, grouped by subjects (L, left and R, right). The vertices of the solid shape within each heptagon show the percent activation of each of the seven pathways. A shape with a larger area generally signifies greater overall pathway activation for that DBS lead.

### Relationship between changes in gait and pathway activation

In considering the pathway activation maps (**Fig. 3**), three of the 10 subjects (Subjects 5-7) had similar pathway activations, characterized by robust bilateral activation of the PPN→GP pathway, GPi→PPN pathway, and the hyperdirect pathway, and with less activation in the other pathways. DBS through either lead in these subjects resulted a primarily shortened (worsening of) step length (**Fig. 2**). The DBS leads with the largest percent increase in step length (Subject 3 left/right and Subject 10 left), showed relatively less activation of the hyperdirect pathway while still activating the PPN→GP pathway robustly. To further investigate these findings, the steady-state changes in step length for each DBS lead tested and the corresponding pathway activation for that DBS lead were incorporated into a linear mixed effect model (**Fig. 4**). In the model, positive coefficients corresponded to increases (improvement) in step length and negative coefficients corresponded to a shortened step length. Coefficient estimates showed that the PPN→GP pathway had statistically significant positive coefficient (*p*=0.025; F-test, df=32) and GPi→PPN and the hyperdirect pathway had a significant negative coefficient (*p*=0.006 and *p*=0.0024, respectively; F-test, df=32). The STN efferent pathways and the cstIC pathway did not reach statistical significance.

**Figure 4.**
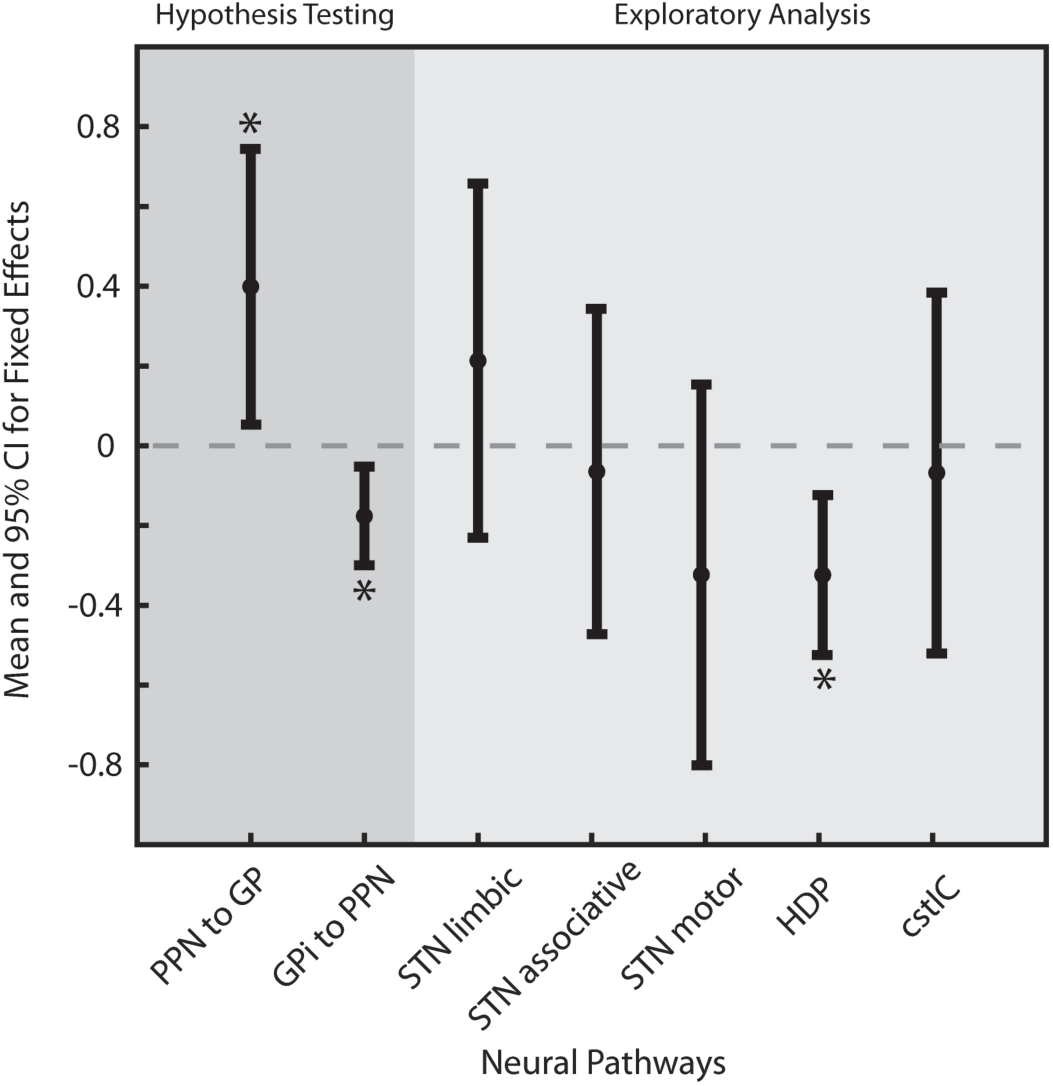
Mixed-effects model results for steady-state step length change over time (including both wash-in and wash-out data from the foot contralateral to the DBS lead). * Statistically significant at *p* <0.05 for an F-test.

## Discussion

Gait disturbances in idiopathic PD are common in the late stages of PD and, when they occur early, predict faster progression and poorer prognosis. Longitudinal studies have also shown that patients initially classified as tremor predominant PD can be reclassified into the postural instability and gait difficulty (PIGD) type as the disease progresses^38–40^. PIGD manifests as a shortened stride length, increased cadence and decreased step length^41^. The responsiveness of PIGD to standard clinical care is limited and STN-DBS has mixed effects on gait measures including step length^42^. The difficulty of optimizing STN-DBS specifically for gait dysfunction stems from two sources: 1) there are long wash-in and wash-out times to affect gait symptoms, making it difficult to optimize DBS during a clinical visit^43^ and 2) it is not clear which neural pathways should be targeted to treat gait dysfunction and should be avoided prevent stimulation-induced worsening of gait.

Here, we used a side-specific gait metric (step length) as the outcome measure of DBS. Although the change of step length reached a steady state after a 30 minute wash-in/wash-out for most of the subjects, our approach of fitting a curve to the gait measurements at different time points enabled us to estimate the effect of DBS at steady state for all subjects, as well as allowing us to use a larger number of measurements in each subject, obtained over one hour, reducing measurement noise. Appropriate statistical analysis also allowed us to account for non-independence of the measurements of two hemispheres from a single subject.

Computational modeling studies of STN-DBS have also been useful for making associations between neural pathway activations and clinical observations assessing the degree of therapy or the emergence of side effects. Butson and colleagues first identified locations around the STN that aligned with improvement in bradykinesia and rigidity^44^, suggesting that targeting pathways outside the closed field nucleus of the STN correlated with therapy. Other studies have noted that targeting STN efferent neurons in the motor territory aligns with rest tremor control, and activating the pallidothalamic tract can suppress stimulation-induced dyskinesias^13^. Previous studies, however, have only modeled a few pathways around the STN or relied upon pathway models that are normalized to a standard atlas. Here, we constructed subject-specific pathway models and incorporated seven different neural pathways populated with multi-compartment axonal morphologies and associated the effects of DBS on the neural pathway activations with changes in step length. We tested our prospective hypothesis that activating the pedunculopallidal pathway would increase step length and activation of the pallidopenducular pathway, which sends GABAergic efferents to the mesencephalic locomotor region, would reduce step length.

Our statistical and subject-specific computational modeling results showed a strong positive effect of pedunculopallidal pathway activation on step length. Population-level analysis of the pathway activations revealed that the threshold of activation for PPN→GP pathways was low, and the pathway was highly activated (>90%) at clinical settings with STN-DBS for 85% of the hemispheres (17/20 sides). Although PPN→GP pathway activation was relatively high amongst DBS leads tested, there was a distribution of activation. Omitting high leverage data points did not change the statistically significant positive effect.

The PPN has a heterogeneous cellular structure with axons from the glutamatergic and cholinergic neurons sending projections to the basal ganglia amongst other subcortical structures^45,46^. Optogenetic stimulation of the glutamatergic neurons in the PPN during treadmill walking in rodents caused slowing and disruption of locomotion, with prolonged increase in muscle tone^47,48^. Cholinergic neuron stimulation in the PPN similarly results in a slowing of locomotion, though to a smaller extent^47^. Our patient-specific models suggested that high-frequency stimulation of PPN→GP fibers resulted in an increased step length during ambulation. One explanation could be that high frequency DBS is known to create informational lesions^49^ and synaptic depression that would effectively remove abnormal feedback information about step-length versus candence^41^ through reciprocal connections between basal ganglia and mesencephalic locomotor circuits^20,21,47,48^. Additionally, the antidromic activation of those glutamatergic and cholinergic neurons in the PPN may modulate axonal collaterals that could affect a broader cell population within the mesencephalic locomotor region^18^.

The mixed-effect model results suggested that activation of the pallidopeduncular pathway had a negative effect on step length. This is in agreement with previous findings^23,24^ that showed activation of the ventral pole of the GPi can worsen gait and induce hypokinetic movements in dystonia patients and that stimulation in the Forel Fields H2 can worsen gait dysfunction in patients with Parkinson’s disease^4,6^. Moreover, stimulation of the GPi, which contains GABAergic neurons that project to the PPN^50^, can inhibit PPN activity^21,51^ and thus have a direct influence on gait control^52,53^. In contrast, electrophysiological recordings in humans have suggested that STN neurons, as opposed to fibers of passage, may have a relatively limited role on modulation of gait^53^, which also agrees with our findings.

Previous electrophysiological studies have suggested that one therapeutic mechanism of STN-DBS may be antidromic cortical activation of corticospinal tract axon collaterals (HDP)^54^ as demonstrated in preclinical evaluation of rotational behaviors and increased path length in mice^55^. Studies in humans have shown short-latency activation of motor cortex that may result from both HDP activation and direct activation of the cstIC^56–58^. Such activation has shown some relationship to improvement in clinical ratings or choice of electrode configuration for programming^56,59,57^; however, previous studies have not specifically investigated how HDP activation affects gait. We found that HDP activation was associated with shortened step length. This may be because HDP axons project into the STN as collaterals from cstIC fibers^17^, thus antidromic activation of HDP axons may antidromically affect motor cortex^60^ but also orthodromically affect lower brainstem and spinal cord circuits^17^ with the latter effect causing impaired ability to transmit information related to step length during walking. Direct activation of the cstIC pathway was marginal across all DBS leads tested (**Fig. S2**) as expected, since clinical settings tried to avoid inducing involuntary muscle contractions^61–63^. Lack of significant cstIC activation may explain the lack of significant association between cstIC pathway activation and step length.

Although our study has strengths, it is not without its limitations. The regression analysis relied on a linear approximation of the relationship between pathway activation and step length^63^. Additionally, the pathway activation estimates were based on a single clinically optimized setting in each of the 10 subjects (20 DBS leads), which limited the variance of pathway activation. While we modeled seven unique pathways, there are other electrically excitable structures in and around the STN. For example, GPi neurons coursing through the lenticular fasciculus branch to the centromedian and parafasicular nuclei of thalamus^19^, and these axonal branches were not explicitly accounted for in our models. The positive effect of activation of the PPN→GP pathway could also reflect activation of some other, unmodeled neural process whose activation correlated with activation of the PPN→GP pathway. Further, while the absolute modeled activation percentage depends on a broad range of biophysical model parameters including fiber diameter, our results were based on relative pathway activation, which mitigates this limitation. Our gait measurements also had limitations. Although we attempted to blind subjects to stimulation condition (on/off), this proved impossible in most cases, due to return of non-gait symptoms such as tremor, in the off-stim condition. However, our use of quantitative gait metrics, the consistency of repeated measurements over time, and the fact that the gait effect of stimulation was not always in the expected direction (i.e. sometimes negative, and sometimes opposite on the two sides) suggest that this did not affect our results.

## Authorship Roles

Design (MG, MDJ, and SEC), execution (MG, CL, MS, RP, TP, JK, NH, MDJ, and SEC), analysis (MG, CL, MDJ, and SEC), writing (MG, CL, MDJ, and SEC), and editing of the final version of the manuscript (MG, CL, MS, RP, TP, JK, NH, MDJ, and SEC).

## Acknowledgements

We thank William Guo for help with the DBS lead segmentations. We also thank the Minnesota Supercomputing Institute (MSI) for computational resources to perform the pathway activation models.

## Supplementary Document

### Finite element modeling (FEM)

The imaging data sets enabled constructing subject-specific finite element models of the electrode-tissue interface surrounding each DBS lead^61,64,15,12–14^. The distribution of stimulus-induced tissue potentials was modeled in COMSOL Multiphysics (v5.4). The FEM construction included the following processes: 1) DBS lead localization using postoperative CT scans co-registered to DTI images. 2) Brain segmentation using the BET tool in FSL and then brain-masked T1-weighted image fed into the FSL FAST tool to segment the brain into white matter (WM), grey matter (GM) and cerebrospinal fluid (CSF). Conductivity tensors for each segmented class were calculated using DTI^65^. These were normalized by the isotropic electrical conductivities derived from the median frequency of the applied stimulus, defined by frequencies of 4294 Hz (Medtronic stimulator) and 3049 Hz (Abbott and Boston Scientific stimulators). (CSF: σ=2.0 S/m, ε_r_=109; 4924 Hz: WM: σ=0.065 S/m, ε_r_=23361; GM: σ=0.11 S/m, ε_r_=48617; 3049 Hz: WM: σ=0.065 S/m, ε_r_=29790; GM: σ=0.11 S/m, ε_r_=65898). A 0.25 mm encapsulation layer around the DBS lead was modeled with isotropic electrical properties of white matter. Lumped-tissue surrounding the brain was modeled with isotropic electrical properties of σ = 0.23 S/m and ε_r_ = 1 in each subject’s brain model^64^.

A multi-resolution FEM mesh was generated using Delaunay triangulation. The tetrahedral element spatial resolution was 0.07 – 0.2 mm near the DBS lead and up to 20 mm at the surface of the scalp. The resulting volumes were meshed quadratic tetrahedral elements. For monopolar DBS settings, the base of the neck was assigned as ground (i.e. 0 V, Dirichlet boundary condition), while the rest of the scalp surface and the DBS lead shaft were assigned zero current density (i.e. Neumann boundary condition). To estimate the spatial distribution of electric potential, simulations were run at a single AC frequency, corresponding to the median frequency of the stimulus waveform as described above.

### Multi-compartment NEURON Models

Neurons within the STN and axons of passage were populated with multi-compartment morphologies constructed in MATLAB^12–14^. The simulated axons of passage were built as myelinated cable models^66,67^ based on either an axon diameter of 0.8 μm for the HDP or 1.6 μm for all other pathways. The node diameter was 0.7 μm (HDP) and 1.4μm (other fiber pathways). Number of lamellae was 20 (HDP) and 30 (other fiber pathways). Additionally, passive nodes were instantiated at first and last nodes to prevent low-threshold terminal activation.

### Modeling of DBS lead tissue displacement

We also incorporated spatial remapping of the STN neurons and axonal fibers of passage to account for tissue deformation that can occur around a DBS lead following implantation^68–70^. The three-dimensional coordinates for each compartment of both STN neurons and fibers of passage were displaced using the following formula: r + R*exp(-r), where “r” is the distance from the closest compartment to the center of the DBS lead and R is the sum of the radius of the DBS lead (635 um) and thickness of the encapsulation layer (250 um). For the STN neuron models, these morphological templates were translated based on the above equation, whereas for the axonal fibers of passage, the translation was performed as part of generating the axonal compartment morphologies. This remapping resulted in neural processes near to (distant from) the DBS lead having large (small) displacements. We also defined the DBS lead as an exclusion surface and modeled the neural processes such that they did not intersect with this surface.

---

**Supplemental Figure 1.**
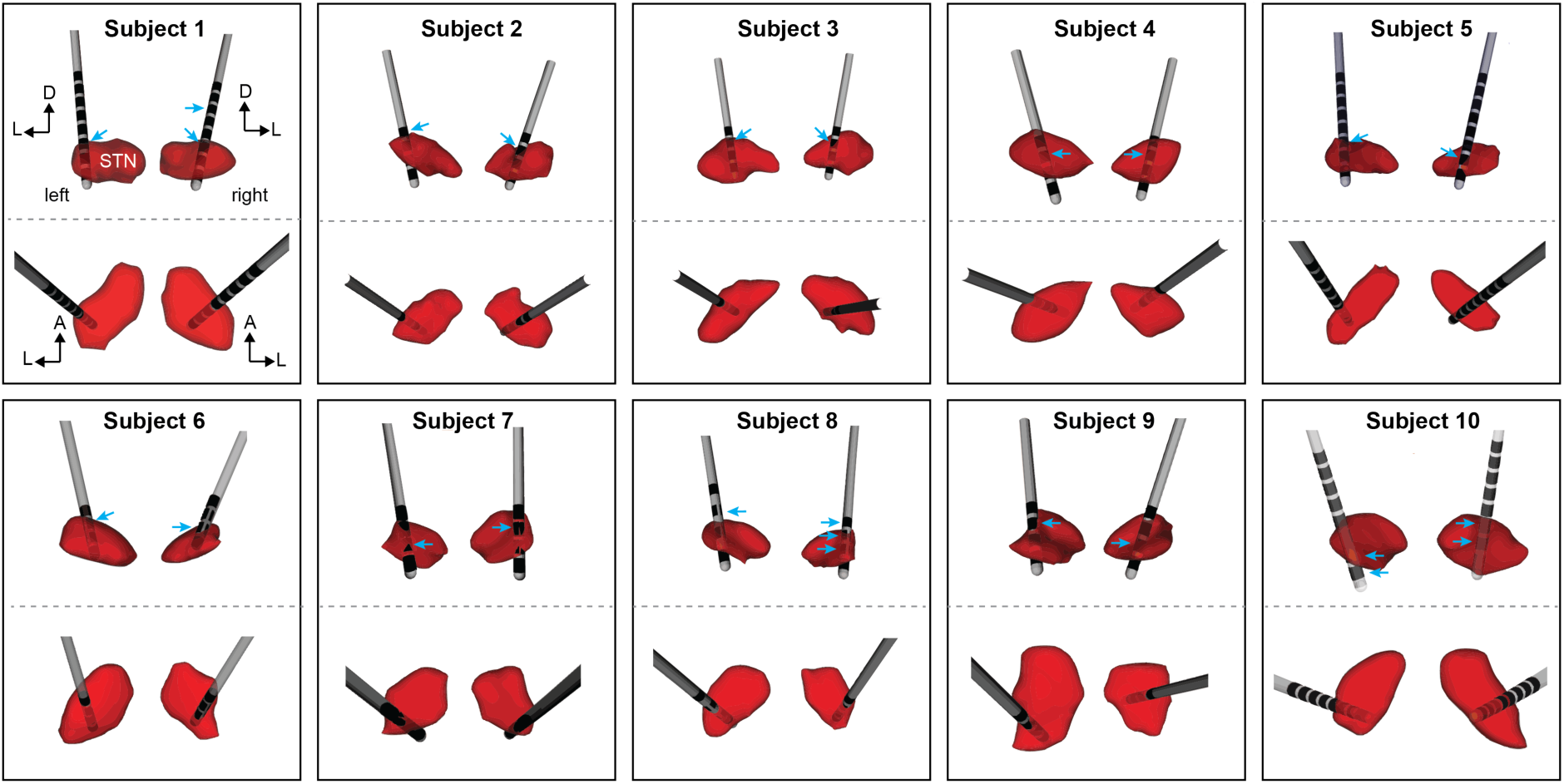
Bilateral DBS lead placement and orientation within the STNs of each subject. Top views show the coronal perspective and the bottom views show the axial perspective. Cyan arrows signify the active electrodes (see Figure 3 for stimulation settings).

**Supplemental Figure 2.**
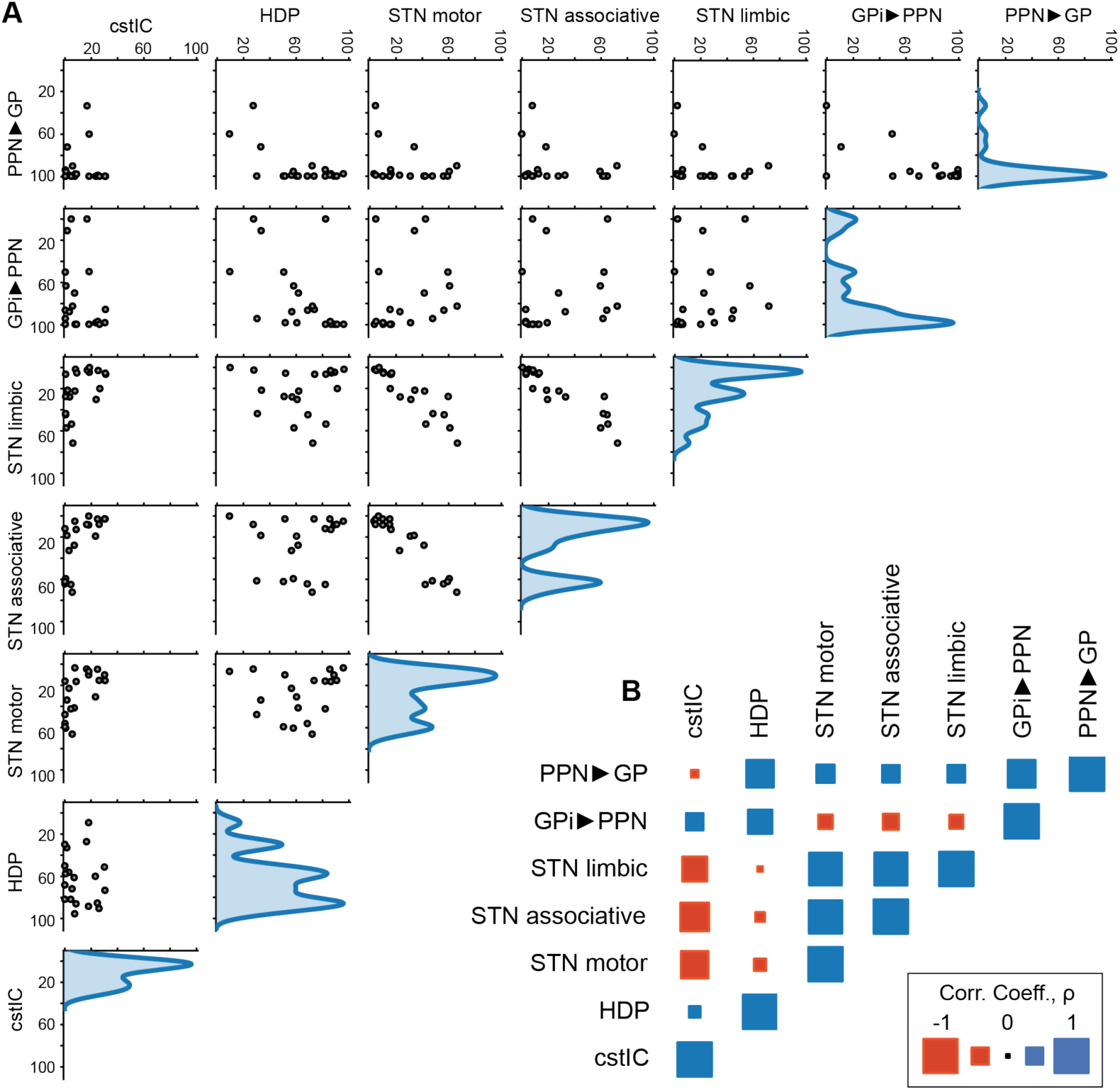
Pathway activation profiles for clinically optimized stimulation settings across twenty DBS leads. Population-level analysis of the similarity in activation profiles between modeled pathways as shown through **(A)** scatterplots and **(B)** correlation analysis.

